# The FAIR Data Point Populator: collaborative FAIRification and population of FAIR Data Points

**DOI:** 10.1101/2024.09.06.611505

**Authors:** Daphne Wijnbergen, Rajaram Kaliyaperumal, Kees Burger, Luiz Olavo Bonino da Silva Santos, Barend Mons, Marco Roos, Eleni Mina

## Abstract

**Background:** Use of the FAIR principles (Findable, Accessible, Interoperable and Reusable) allows the rapidly growing number of biomedical datasets to be optimally (re)used. An important aspect of the FAIR principles is metadata. The FAIR Data Point specifications and reference implementation have been designed as an example on how to publish metadata according to the FAIR principles. Various tools to create metadata have been created, but many of these have limitations, such as interfaces that are not intuitive, metadata that does not adhere to a common metadata schema, limited scalability, and inefficient collaboration. We aim to address these limitations in the FAIR Data Point Populator.

**Results:** The FAIR Data Point Populator consists of a GitHub workflow together with Excel templates that have tooltips, validation and documentation. The Excel templates are targeted towards non-technical users, and can be used collaboratively in online spreadsheet software. A more technical user then uses the GitHub workflow to read multiple entries in the Excel sheets, and transform it into machine readable metadata. This metadata is then automatically uploaded to a connected FAIR Data Point. We applied the FAIR Data Point Populator on the metadata of two datasets, and a patient registry. We were then able to run a query on the FAIR Data Point Index, in order to retrieve one of the datasets.

**Conclusion:** The FAIR Data Point Populator addresses several limitations of other tools. It makes creating metadata easier, ensures adherence to a common metadata schema, allows bulk creation of metadata entries and increases collaboration. As a result of this, the barrier of entry for FAIRification is lower, which enables the creation of FAIR data by more people.

## Background

The rate of data acquisition in biomedical research is rapidly increasing [1]. In order to enable optimal (re)use of this growing number of datasets, it is important that they are Findable, Accessible, Interoperable and Reusable (FAIR) for both humans and machines [2]. An important aspect of increasing the degree by which data is FAIR (FAIRification), is the provisioning of rich metadata (Principle R1).

The FAIR Data Point (FDP) reference application [3] has been designed to serve as an example of how to publish metadata according to the FAIR principles. The FDP specification consists of a metadata schema structure based on the Data Catalog Vocabulary (DCAT), which has been designed to share metadata of different types of digital objects in an interoperable manner [4]. The FDP reference application serves as a reference implementation of the FDP specification [5]. It has interfaces (webpages, documents, API and SPARQL endpoint) that present the metadata following this schema, and allows for creation and modification of entries both for humans and machines. The machine readable metadata for the FDP is represented as RDF (Resource Description Framework) documents [6] in order to adhere to the FAIR principle I1, of formal knowledge representation of metadata. Finally, there is service for FDPs that indexes metadata from other FAIR Data Points, and allows queries to be executed on all this metadata simultaneously.

Various tools have been developed for creating and publishing metadata. These include tools such as CEDAR [7], RightField [8], the FAIR data station [9], the OpenRefine metadata extension [10] and the FAIR metadata editor [11]. These tools all have their own design philosophy, each with their own advantages and disadvantages. For example, CEDAR, OpenRefine and the FAIR metadata editor work primarily with web interfaces, while RightField and the FAIR data station are focused on Excel. Some of the tools are more adaptable, while some of the tools are more easy to use.

There are some limitations to existing tools. Most importantly, many tools are not easy to use for people who are not familiar with FAIR. They also lack a convenient way to make sure that the metadata is published and indexed, which is one of the requirements for the FAIR principles. Further, many tools require the need to add every entry manually, are not easy to use in collaborations and do not offer version control of the metadata. On the side of availability, many tools require local installations, or are only available online, and thus rely on either local installations, or the continued hosting of the provider of the tool. Finally, some tools create metadata that do not adhere to a common metadata schema.

In this work, we implement a FAIRification tool called the “FAIR Data Point Populator”, that aims to address these limitations, and hereby make the provisioning of FAIR metadata as easy as possible. The tool is written in Python, and is accessible as either a GitHub workflow, or a Jupyter notebook. It reads an Excel template, and automatically publishes the contained metadata in a FDP application as RDF.

## Implementation

The FDP Populator consists of an Excel template together with a python tool. The Excel template contains the metadata that is needed to populate the FDP, which is based on the DCAT schema. There are two sheets in the Excel template that can be filled in, the dataset sheet, and the distribution sheet. The column names in these sheets are shown in table 1. In order to aid in filling in the template, all of the columns are described in the readme file in the same GitHub repository as the template. In addition, the template contains tooltips to describe what information should be filled in every column, and what the requirements are. There is also validation applied on the Excel fields to give warning when metadata does not meet the requirements. For example, when a link is not actually a link. Some extra metadata, like the date of publication and modification, are automatically added by the FDP.

**Table 1:**
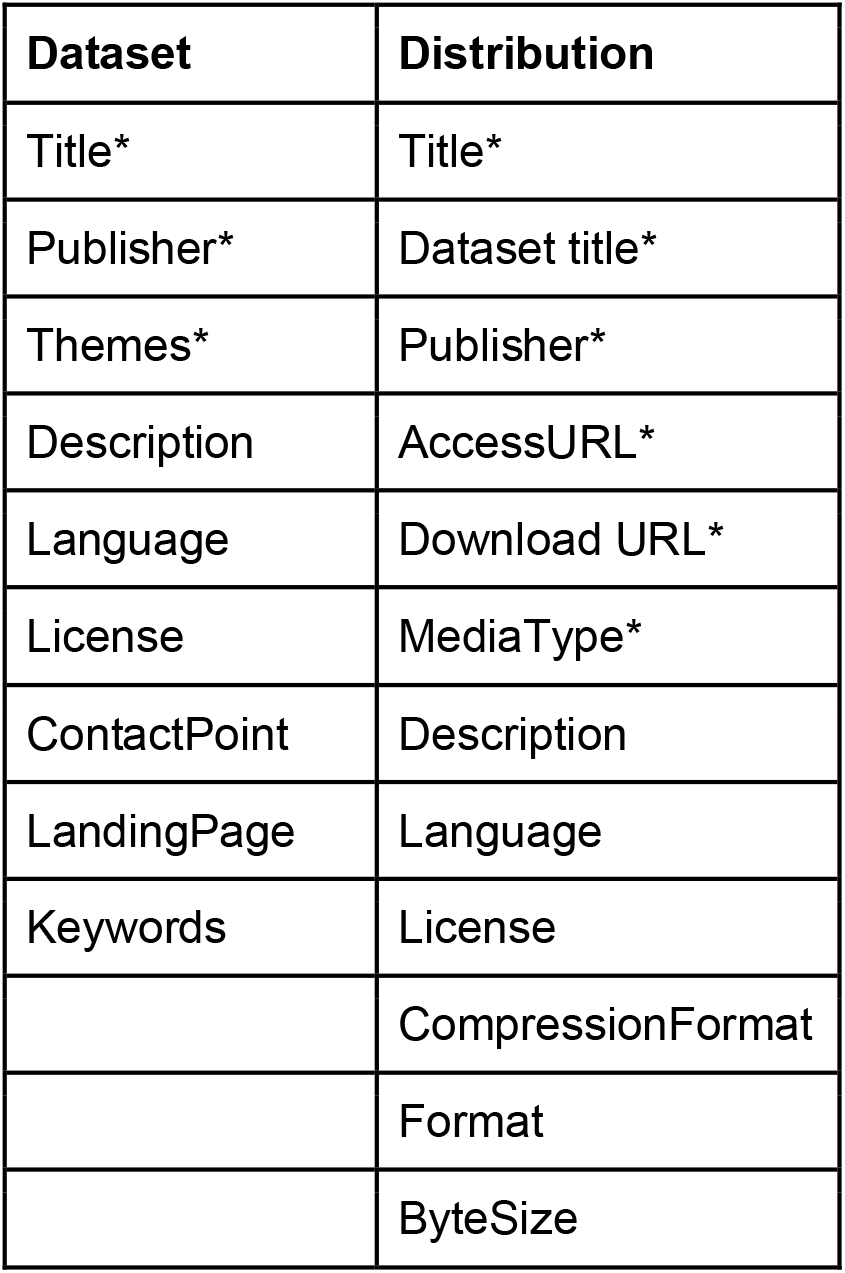
The column names in the Excel template for the “dataset” sheet, and the “distribution” sheets. Required columns are marked with an asterisk.

| **Dataset** | **Distribution** |
| --- | --- |
| Title* | Title* |
| Publisher* | Dataset title* |
| Themes* | Publisher* |
| Description | AccessURL* |
| Language | Download URL* |
| License | MediaType* |
| ContactPoint | Description |
| LandingPage | Language |
| Keywords | License |
|  | CompressionFormat |
|  | Format |
|  | ByteSize |

This Excel template is then used by the FDP Populator tool, which performs the actual publication process. A diagram of its function is shown in figure 1. The tool reads the Excel file, and turns the information into RDF which follows the FDP metadata schema. Finally, the RDF is automatically sent to a FDP using the API of the FDP. URIs of the newly made resources are used in the RDF of the succeeding resources. Both the conversion and the publishing scripts are written in python. We chose to implement these steps as a GitHub workflow in order to make use of GitHubs features. A Jupyter notebook that runs the FDP Populator has also been created as an alternative that does not depend on GitHub.

**Figure 1:**
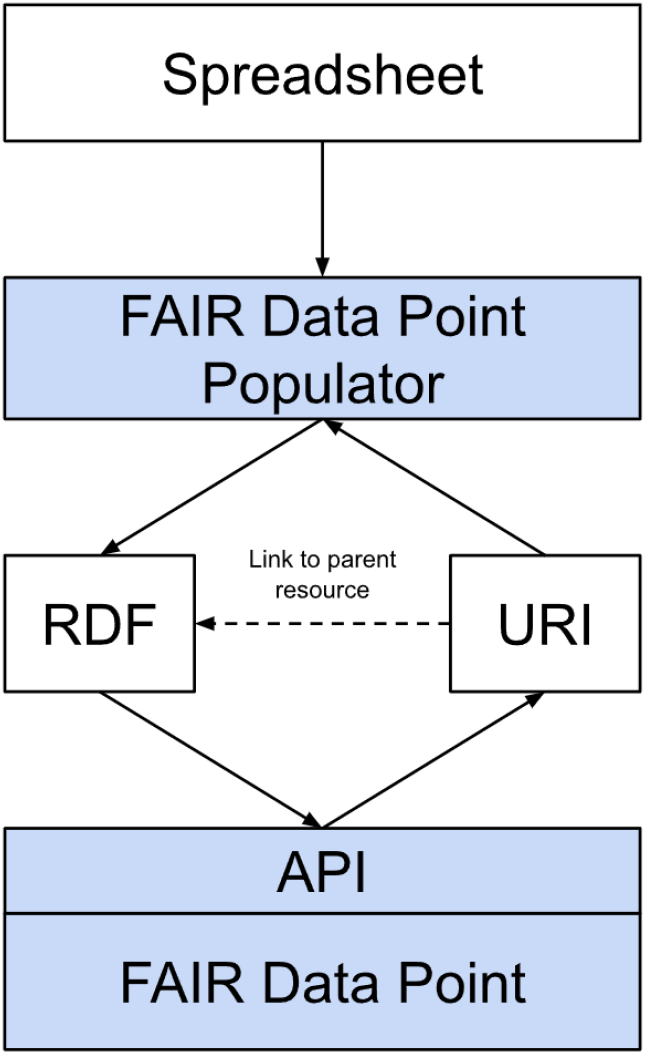
The workflow of the FAIR Data Point Populator. A spreadsheet is read into the FPD Populator, then converted to RDF. The RDF is sent to the FDP after which the FDP returns a URI to this new entry. The FDP Populator can then link to this URI from the following resources.

In order to make things as easy as possible for data owners, the tool has been developed with a collaboration with a data steward in mind. In this case, the data owners have the role of filling in the Excel template, with metadata, while the data steward has the role of uploading this template to GitHub, and starting the workflow that runs the FDP Populator. A diagram of this workflow is shown in figure 2.

**Figure 2:**
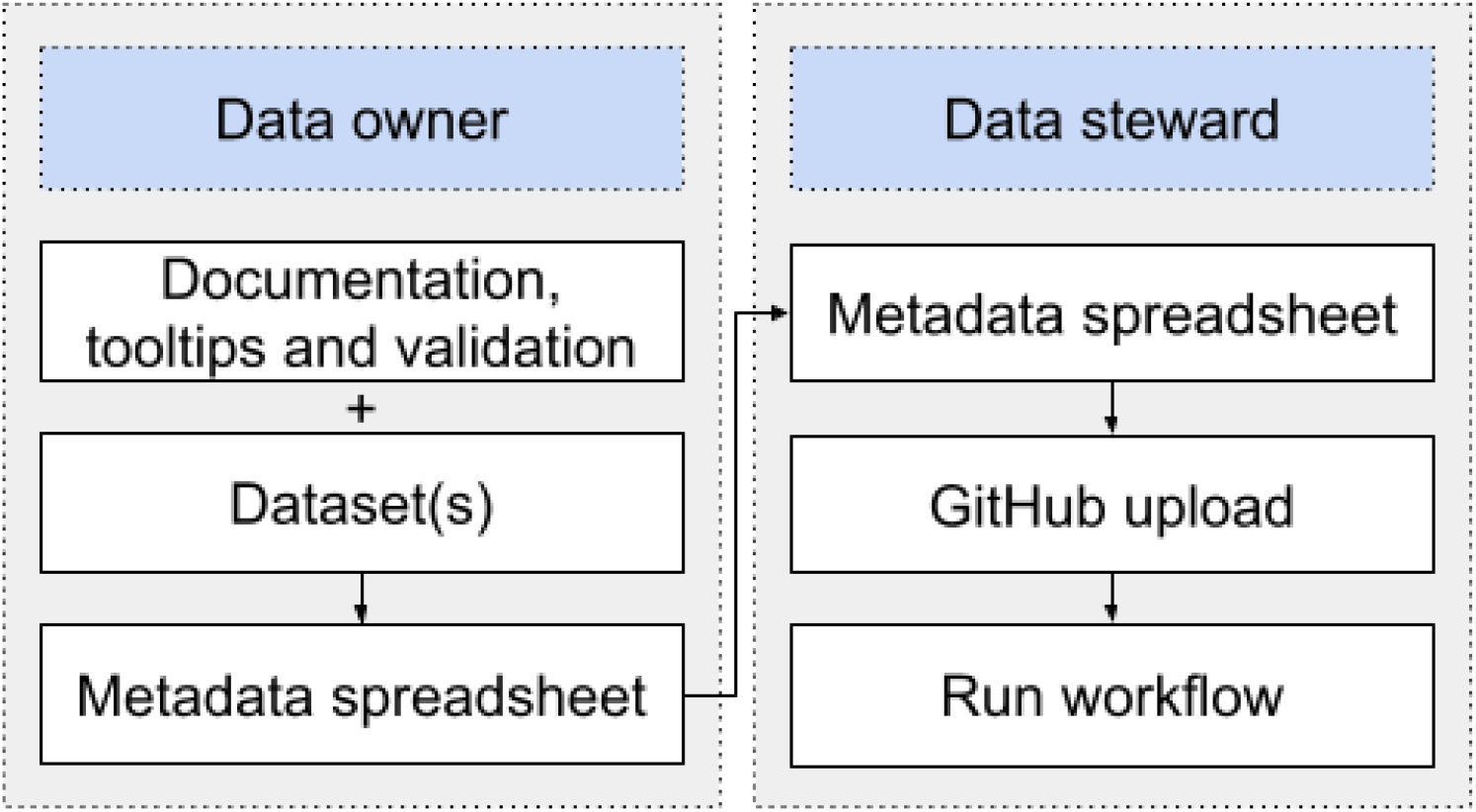
The workflow for users, split into data owner and data steward. The data owner creates the metadata spreadsheet for their data with help from documentation tooltips and validation. The data steward uploads this to GitHub, and runs the GitHub workflow.

The European Joint Programme on Rare Diseases (EJP RD) has created a metadata schema to describe rare disease resources [12]. This schema is focused on making it possible to find resources such as biobanks, patient registries and guidelines. Like the FDP specification, The EJP RD metadata schema is an extension of the DCAT schema. We extended the FDP Populator to use the Excel template from the EJP RD and thereby help disease researchers to describe and publish their resources in a FDP with the help of the FDP populator.

## Results

### Collaborative FAIRification of datasets using automated publication

To show the application of the FDP Populator, we applied it on a number of resources in the context of the EJP RD consortium. First, we applied it on a collection of Inclusion Body Myositis data. These datasets included raw and normalized transcriptomics, raw and normalized miRNA transcriptomics, and whole exome sequencing together with sample information. All of the metadata for these entries were filled in the Excel template in collaboration with the data owners. The dataset sheet with these entries is shown in figure 3. We then ran the FDP Populator on this Excel file in order to generate the corresponding RDF, which can be seen in listing 1 and 2. Finally, immediately after the last step, the tool also automatically populated the FDP with this RDF, which can be seen in figure 4. This made it unnecessary to upload all the metadata manually. Also note that the datasets were all published in one run of the tool, without the need to repeat the process for every dataset. The input Excel file and output RDF can be found at: https://github.com/jdwijnbergen/fdp-populator-usecases, along with a link of the resulting entries in the FDP.

**Figure 3:**
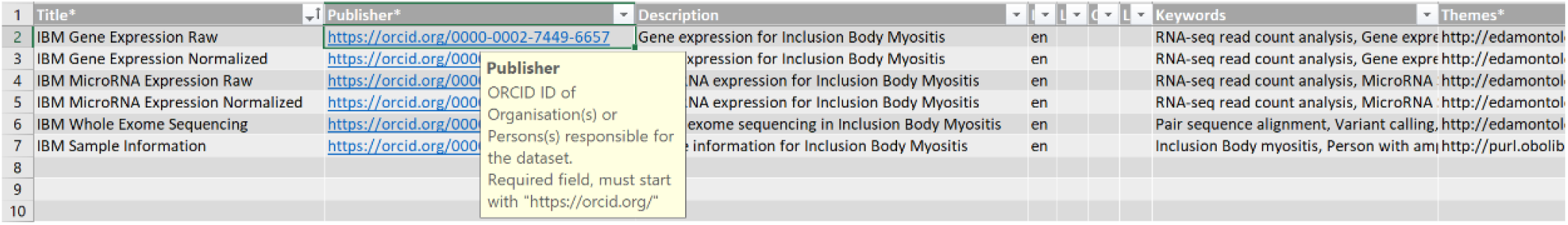
Part of the dataset sheet of the Excel template filled in with metadata for the Inclusion Body myositis datasets. The tooltip for the publisher column is shown. The template also contains validation. Required fields are marked with an asterisk, and have been filled. Some of the optional fields have been left empty.

**Figure 4:**
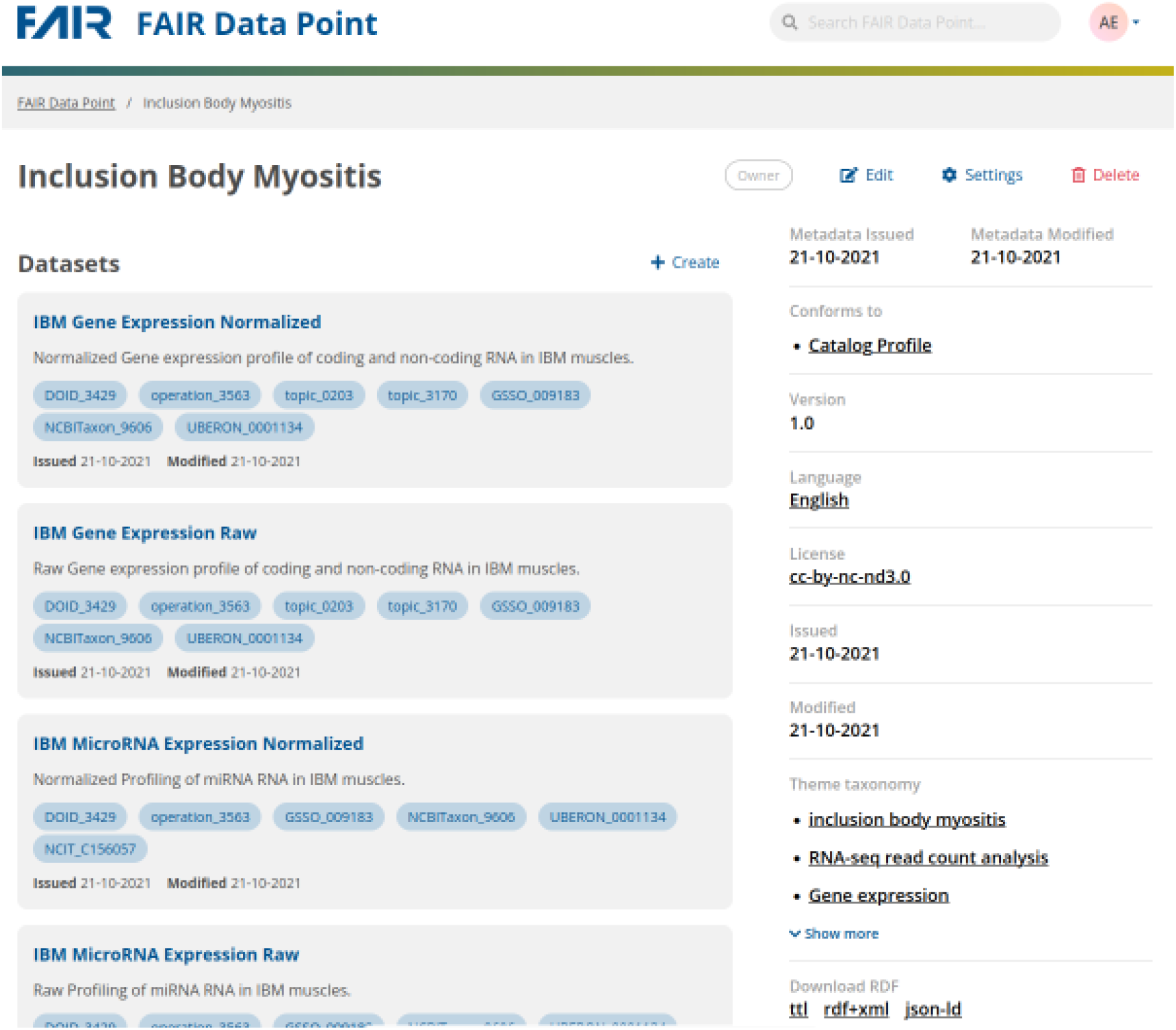
The metadata after it is automatically published on the FAIR Data Point. The dataset metadata can be browsed further, and data downloaded where it is available.

We also show the application of the FDP Populator in two other use cases. The first one is three datasets for CAKUT (Congenital Anomalies of the Kidney and Urinary Tract). This included a mirnome, peptidome and proteome dataset. In this use case, we took advantage of online spreadsheet software to fill in the Excel template. We found that users were able to work on different sections of the Excel template simultaneously, which greatly sped up the process. The resulting Excel file and RDF can be found in the same GitHub repository, along with the link to the entries on the FDP. Finally, showing the implementation of the EJP RD metadata schema in our tool, we also applied the tool on the ERKNet. ERKNet is a patient registry that focuses on rare kidney diseases. In this case, we filled in the EJP RD Excel template, which is adapted to the EJP RD metadata schema. The Excel file and the corresponding output can be found in the same GitHub repository.

### The FAIRified resources are findable through RDF queries

In order to find datasets related to CAKUT, and show that retrieving these datasets is possible for humans and machines, we created a SPARQL (SPARQL Protocol and RDF Query Language) query to run on RDF. This query retrieves entities that are an instance of the dataset class in the DCAT schema, and have CAKUT (in ontologized form) as one of their themes. The query is shown in figure 5. This query was executed on the FAIR Data Point index on the 7th of August 2023. At this time, there were 46 active FAIR Data Points, including the FDP with the CAKUT metadata, on which the index applied the query. We were able to find the three CAKUT datasets that we applied the FDP Populator on (figure 5). It is important to note that this query is correct not only for metadata that is created by our tool, but for any metadata that conforms to the DCAT schema.

**Figure 5:**
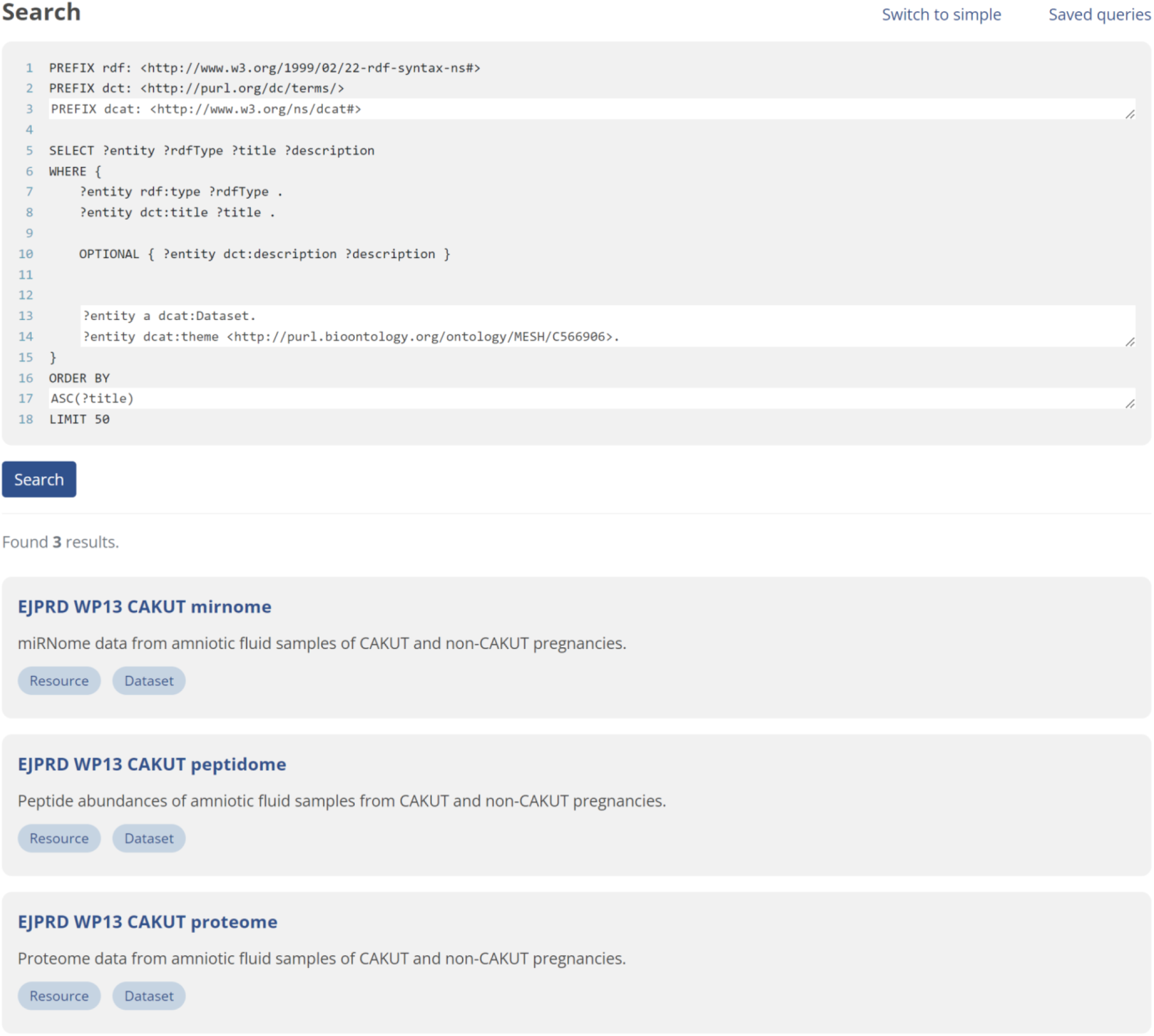
The results of a SPARQL query that looks for any datasets that have CAKUT as their theme. The FAIRified CAKUT datasets are found in the results.

## Discussion

In this work, we created the FDP Populator, which addresses some limitations of previous tools to make it easy to use, enable collaboration, and ensure metadata adhering to a common metadata schema among other advantages. We applied the FDP Populator on a number of use cases, and show that the created metadata allows the data to be found by humans and machines.

The FDP Populator has four main advantages. Firstly, the FPD Populator primary objective is to simplify the provisioning of FAIR metadata, and hereby lower the barrier of entry. This is achieved by letting the data owners use the Excel template to describe their resources, which is likely done in software that they are familiar with. Secondly, the FDP Populator allows for the bulk upload of multiple entries to a catalog. This makes the creation of a large number of entries in a catalog feasible. Thirdly, the template can be filled in simultaneously by multiple people, which reduces the time needed for FAIRification. Fourthly, the tool can be run as a GitHub workflow, or as a Python notebook, which reduces the reliance on a local or hosted tool. Additionally, as a GitHub workflow, it keeps the version history to ensure no metadata is lost.

The FAIR Data Populator also improves the interoperability of the created metadata. Many tools are very flexible, and allow any type of triples to be made. This however can result in RDF that is not easy to use and interpret. The FAIR Data Populator instead adheres to the commonly used DCAT metadata schema as it is implemented in the FDP, and in the EJP RD extension hereof, which ensures interoperability with other resources.

The FDP Populator also has some limitations. It is relatively inflexible in regards to data schemas due to the focus on the FDP and EJP RD metadata schemas. This is a tradeoff made to ensure it outputs proper metadata, though it can be limiting for some use cases. The FDP Populator could be extended to create Excel templates based on FDP metadata schemas, in order to work with rich domain specific metadata schemas. This would however also decrease the user friendliness, as documentation, tooltips and validation would have to be created by the administrator, and a schema would have to be available on a FDP. Further, the FDP Populator also relies on the access to a FAIR Data Point, since it does not host the metadata itself, so a FAIR Data Point needs to be created. Finally, the tool is not hosted online as a webtool, which would lower the barrier of entry even more.

## Conclusion

In this work we created the FAIR Data Populator with the goal of addressing limitations of current tools in the field. Our tool creates RDF from information filled in an Excel template, and automatically uploads this to a connected FDP. It improves the ease of use through using an Excel template with documentation, automatic publication of results, allowing the bulk creation of FDP entries within a catalog, and allowing the use of collaborative software such as online spreadsheet software and GitHub, and. It is less reliant on hosting by external parties, and if used as a GitHub tool, keeps track of the version history of the input metadata. Finally, it also improves the operability of the metadata by adhering to commonly used specifications like the FDP specification, and the EJP RD metadata schema which are based on the DCAT schema.

With these advantages, the barrier of entry of FAIRification will be lower. Together with the use of commonly used data schemas, this ensures that more data will be FAIR.

## Availability and requirements

**Project name:** FAIR Data Point Populator

**Project home page:** https://github.com/jdwijnbergen/fdp-populator

**Operating system(s):** Platform independent

**Programming language:** Python

**Other requirements:** GitHub or Google Colab. For local use, Python3 with required packages.

**License:** MIT

**Any restrictions to use by non-academics:** no

## List of abbreviations

CAKUT: Congenital Anomalies of the Kidney and Urinary Tract
DCAT: Data Catalog Vocabulary
EJP RD: European Joint Programme on Rare Diseases
FAIR: Findable, Accessible, Interoperable and Reusable
FDP: FAIR Data Point
RDF: Resource Description Framework
SPARQL: SPARQL Protocol and RDF Query Language

## Availability of data and materials

The FDP Template is available at https://github.com/LUMC-BioSemantics/EJP-RD-WP13-FDP-template, The EJP RD template is available at https://github.com/ejp-rd-vp/resource-metadata-schema/tree/master/template and the repository with use cases is available at https://github.com/jdwijnbergen/fdp-populator-usecases.

## Authors’ contribution

D.W. and R.K.: Conceptualization, development and application on use cases. K.B.: Infrastructure support. D.W. and E.M.: Manuscript preparation. E.M. and M.R.: Supervision. D.W., R.K., K.B. L.S., B.M., M.R. and E.M., contributed to the manuscript, reviewed the manuscript, and gave approval for publication.

## Acknowledgements

We would like to thank Henriette for the creation of the EJP RD metadata schema, Mirdul Johari for the collaboration on the Inclusion Body Myositis use case, Friederike Ehrhart for the collaboration on the CAKUT use case, and Jose Ramírez García for the collaboration on the ERKNet use case.

---

**Listing 1:**
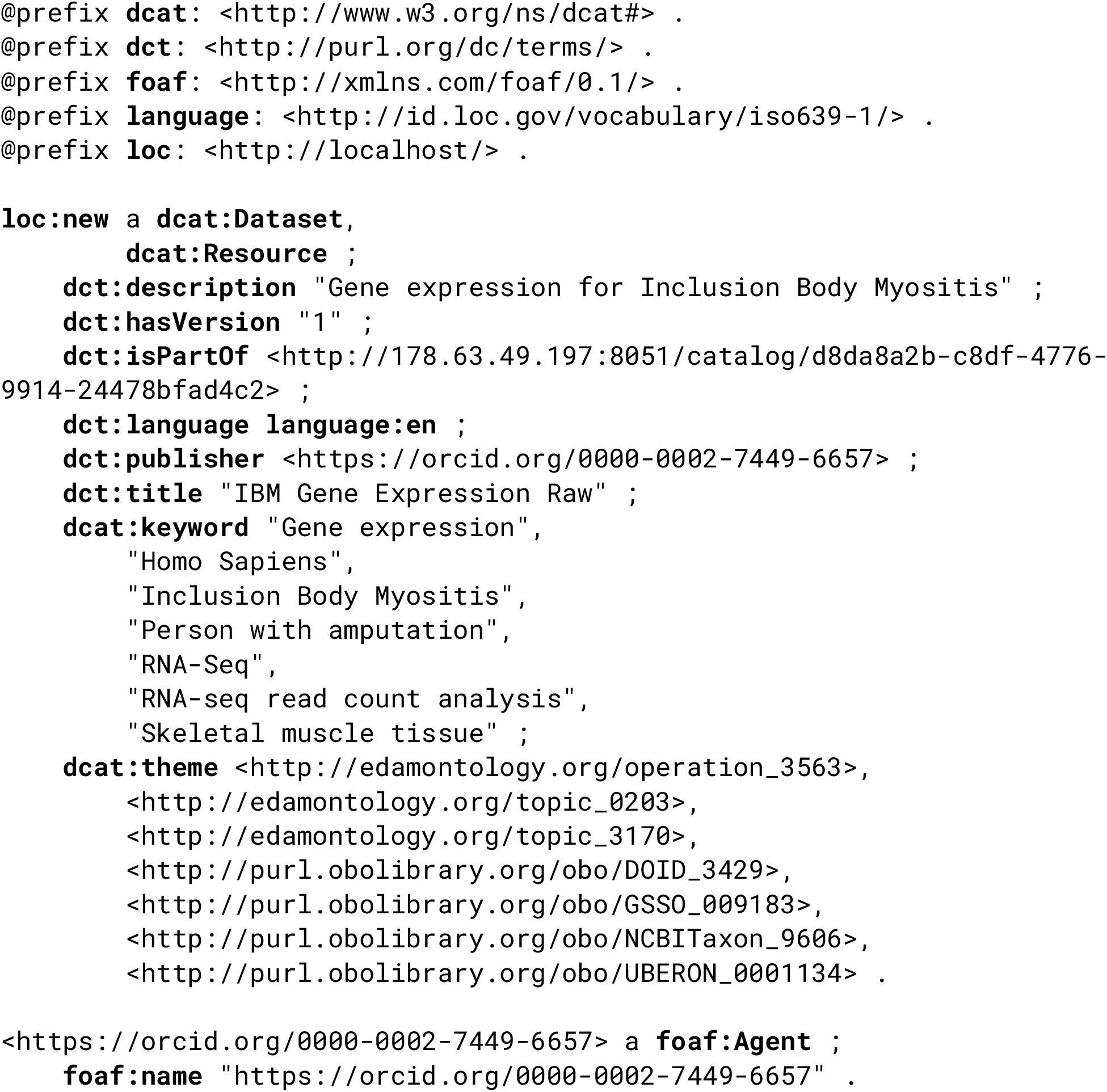
The RDF metadata created for the raw expression IBM dataset expressed in the Turtle RDF format. Note that the FDP adds additional triples to this.

**Listing 2:**
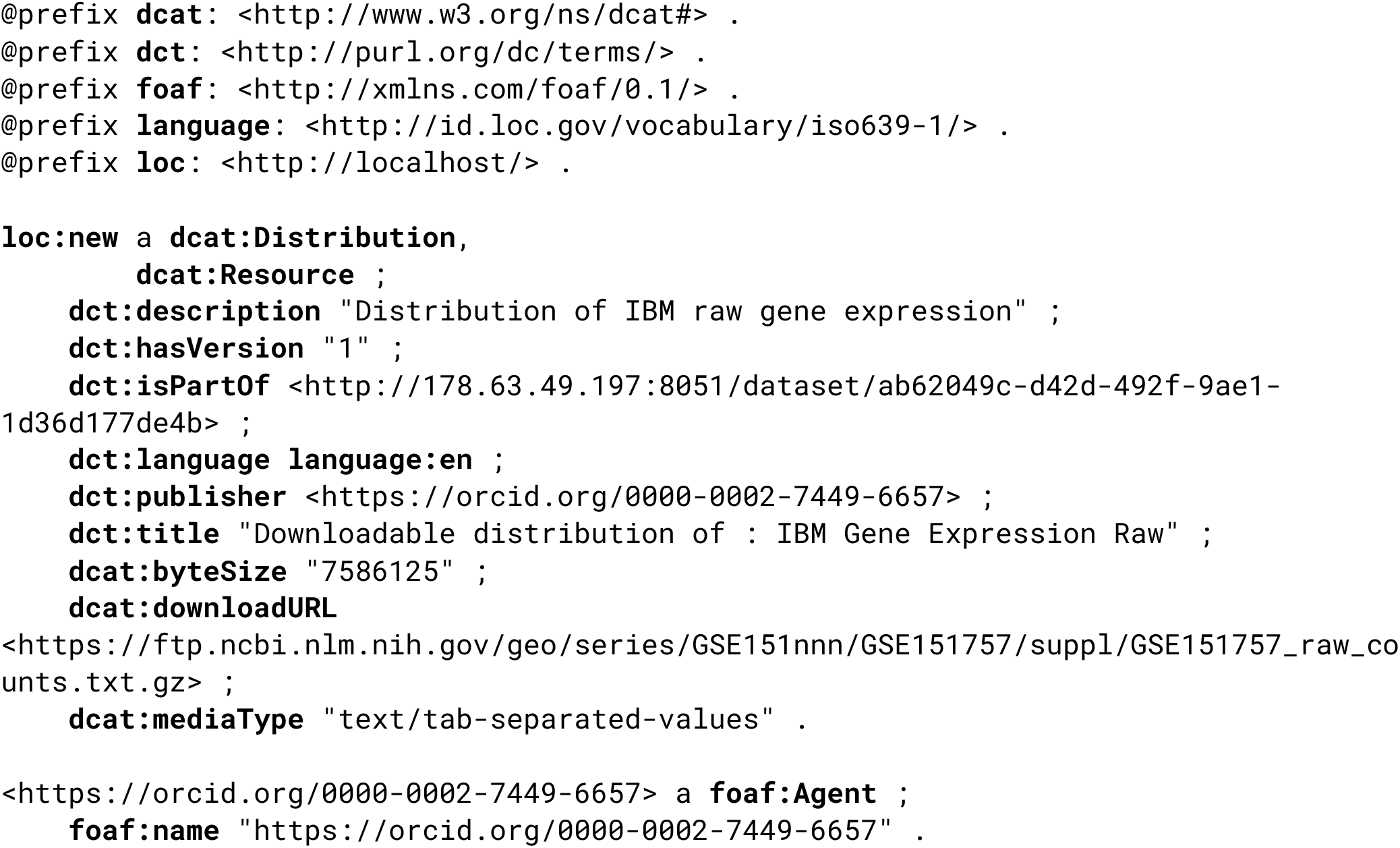
The RDF metadata created for the raw expression IBM distribution expressed in the Turtle RDF format.. Note that the FDP adds additional triples to this.

